# MMD-DTA: A multi-modal deep learning framework for drug-target binding affinity and binding region prediction

**DOI:** 10.1101/2023.09.19.558555

**Authors:** Liwei Liu, Qi Zhang, Yuxiao Wei, Shengli Zhang, Bo Liao

## Abstract

The prediction of drug-target affinity (DTA) plays an important role in the development of drugs and the discovery of potential drug targets. In recent years, computer-assisted DTA prediction has become an important method in this field. In this work, we propose a multi-modal deep learning framework for drug-target binding affinity and binding region prediction, namely MMD-DTA. The model can predict DTA while unsupervised learning of drug-target binding regions. The experimental results show that MMD-DTA performs better than the existing models on the main evaluation metrics. In addition, external validation results show that MMD-DTA improves the generalization ability of the model by integrating sequence information and structural information of drugs and targets, and the model trained on the benchmark dataset can be well generalized to independent virtual screening tasks. Visualization of drug-target binding region prediction shows the powerful interpretability of MMD-DTA, which has important implications for exploring the functional regions of drug molecules acting on proteins.

## 1. Introduction

The prediction of drug-target affinity (DTA) plays an important role in drug development and the discovery of potential drug targets. However, traditional biological experiments to verify DTA are a costly, inefficient, and time-consuming process [1–2]. The results of the survey indicate that the economic cost of developing a new drug is approximately $2.6 billion and the time cost is 17 years [3–5]. In recent years, computer-assisted drug-target interaction (DTI) prediction has become an important method in this field. However, the DTI prediction problem is often regarded as a binary classification problem, which seriously affects our ability to quantitatively reflect the strength of the interaction between drug targets. Binding affinity (binding free energy) is used to express the affinity of a given ligand to a particular target and is invaluable information for drug designers. Without this information, interaction data is not sufficient for lead optimization. For example, a ligand can have a binding affinity of –5 kcal/mol, and another ligand can have a much stronger affinity of –11 kcal/mol. Those are different affinities and offer different prospects for optimization. Therefore, the prediction of drug-target affinity (DTA) makes up for part of the deficiency of the DTI prediction.

In 2014, Tang *et al.* for the first time regarded the DTI prediction problem as a regression problem and proposed the famous KIBA dataset [6]. In 2015, Pahikkala *et al.* predicted DTA using similarity scores from drug-target pairs defined by the Kronecker product [7]. In 2017, He *et al.* proposed the concept of DTA prediction and the SimBoost model based on quantile regression to calculate the confidence of DTA prediction [8]. With the wide application of deep learning, Öztürk *et al.* proposed DeepDTA [9] in 2018, which is a method to extract drug and target sequence features based on a convolutional neural network (CNN) and at the same time applying a fully connected neural network (FCNN) for DTA prediction. In 2019, Zhao et al. developed a model called AttentionDTA [10], which uses the attention mechanism to realize the interaction of features and obtain a more effective feature representation. In the same year, Öztürk *et al.* proposed the WideDTA model based on drug and protein sequence features, protein domain features and motifs, and the features of the maximum common substructure words [11]. In 2020, Nguyen *et al.* used the graph convolutional neural network (GCN) to extract the structural information contained in the drug molecule graph and built the GraphDRP model [12], which achieved better prediction accuracy compared to the previous DTA prediction methods that only used sequence features. Subsequently, Jiang *et al.* improved the GraphDRP model by adding structural information from proteins and proposed a DTA prediction model DGraphDTA based on a protein contact map and a graph neural network (GNN) [13]. In 2021, Bahi *et al.* developed a DTA prediction model based on stacked autoencoder (SAE), which improved the accuracy of prediction by using the weight of the initialized CNN [14]. In 2022, Hua *et al.* applied the attention mechanism to extract multi-scale protein features and fingerprint information to construct a CPInformer model to predict compound protein interaction (CPI) [15]. In the same year, Nguyen *et al.* introduced an algorithm based on directed message passing graph neural networks (D-MPNN) [16] and cross-attention networks to predict potential CPI [17]. Recently, in 2023, Hu *et al.* proposed the SAM-DTA model [18], which achieved good DTA prediction performance without obtaining any protein sequence correlation information and only using drugs that interact with proteins to characterize them. He *et al.* combined sequence-based and graph-based methods to propose a node-adaptive DTA prediction model called NHGNN-DTA [19].

With the successful application of deep learning in DTA prediction, more and more deep networks have been established in the study of drug-target binding region (BR) prediction. In 2018, the DeepDTA model proposed by Ozturk *et al.* used an attention module to visualize predictions of drug target BR in test datasets. In 2019, Tsubaki *et al.* developed the CPI-GNN model [20], which represented the different importance of different protein subsequences and drug molecules by assigning different weights to protein subsequences. In 2020, Abbasi *et al.* were inspired by the DeepDTA model to construct the DeepCDA model [21], which is a CPI prediction method based on long short-term memory (LSTM) layers [22], and at the same time applied the two-sided attention mechanism to predict drug-target BR. In the same year, Chen *et al.* introduced a CPI algorithm named TransformerCPI that uses deconvolution to reveal important interaction regions between protein subsequences and compound atoms [23]. In 2022, Hua *et al.* improved the TransformerCPI algorithm by using informer [24] to replace Transformer and built the CPInformer model. In 2023, the NHGNN-DTA model proposed by He *et al.* uses a multi-head self-attention mechanism to show the prediction information of binding sites. Recently, Hua *et al.* combined mean square error (MSE) loss and RWing loss function [25] to propose a multi-functional robust model called MFR-DTA [26]. The model treats drug-target BR prediction as a supervised learning task by Mix-Decoder block and a fully connected layer.

Although the previous methods have made great progress in DTA prediction, there are still some shortcomings. On the one hand, DTA prediction methods based on drug Simplified Molecular Input Line Entry System (SMILES) strings and protein amino acid sequences are overhyped and currently have little impact on computational drug design. The use of protein structural features has further implications. Protein molecules typically contain more than 1,000 heavy atoms. The representation of a one-dimensional (1D) protein sequence is insufficient to capture the 3D structural features of the protein, which may be important factors affecting the affinity of drug targets. Some recent studies are trying to represent and predict the 3D structure of proteins [27–29]. However, direct representation of 3D structure information requires a large number of high-dimensional and sparse matrices as input variables [30–32]. Moreover, the 3D protein structure prediction methods often require higher environment configuration and longer running time, and there are certain errors. These inevitable errors will certainly affect the effectiveness of the DTA prediction. Second, most of the previous methods focus on the feature extraction and affinity prediction of the model, but ignore the importance of feature connection and interaction. The integration of features is usually done through simple concatenation, which is actually very detrimental to capturing the relationship between protein and molecular characterization. On the other hand, the current prediction of drug-target BR mostly uses attention mechanism to highlight important regions in protein subsequences, but these highlighted regions are likely to have no relationship with the biological properties of the protein. Therefore, there is much room for improvement in the prediction of drug-target BR.

To make up for the shortcomings, we propose a multi-modal deep learning framework MMD-DTA to predict DTA and drug-target BR. In our study, we first extract containing drug, target comprehensive features of physical and chemical information and structural information. Specifically, for drugs, we use the open-source software RDKit [33] to extract physicochemical information contained in drug molecules from molecular fingerprints, and use the GNN model to extract structural information of drug molecules from drug molecular graphs. For proteins, we used a 1D convolutional neural network (1D CNN) to extract protein sequence information from amino acid sequence, and a target structural feature extraction network to extract protein structural information from the protein distance map constructed by the PconsC4 model. Second, we construct a new feature fusion module using the attention mechanism. This module helps the model better learn the relationship between features and output the deep fusion features with strong expressiveness, achieving the balance of complementarity and contribution between different features. Finally, we design a feature interaction module, which can simultaneously output the drug-target descriptor for DTA prediction and the interaction strength of drug sequence and target sequence at each site for drug-target BR prediction. We input drug-target descriptors into multi-layer perceptron (MLP) to obtain predicted DTA values. After building the model, we used 5-fold cross-validation (5-fold CV) to evaluate the predictive performance of our model on the Davis and KIBA datasets. Meanwhile, we conduct comparative experiments to demonstrate the superiority of our model and to verify the generalizability of MMD-DTA through external validation. To show the interpretability of our model and the prediction ability of drug-target BR, we visually analyze the prediction results of BR. In general, MMD-DTA is an effective model to predict DTA and drug-target BR.

## 2. Materials and methods

### 2.1 Datasets

To verify the effectiveness of our model and to facilitate comparative analysis with other models, we use two famous benchmark datasets, Davis [34] and KIBA [6]. Furthermore, to evaluate the performance of our model in predicting drug-target BR, we use the sc-PDB dataset [35].

- The Davis dataset contains 30056 interactions between 68 drugs and 442 targets. The kinase dissociation constant *K*_*d*_ indicates the binding affinity, with a value between 5.0 and 10.8.
- The KIBA dataset contains 118254 interactions between 229 drugs and 2111 targets. The KIBA score indicates the binding affinity, with a value between 0.0 and 17.2.
- The sc-PDB dataset is a collection of 16 034 binding sites between 4782 drugs and 6326 targets. The binding sites are extracted from the Protein Data Bank (PDB).

### 2.2 MMD-DTA

In this part, we propose a new multimodal deep learning framework, called MMD-DTA, for drug target binding affinity and BR prediction. The overall framework of MMD-DTA is shown in Figure 1, which consists mainly of four modules: Drug information encoding, Protein information encoding, Feature fusion module, Feature interaction module. The following is a detailed introduction to these four modules.

**Figure 1.**
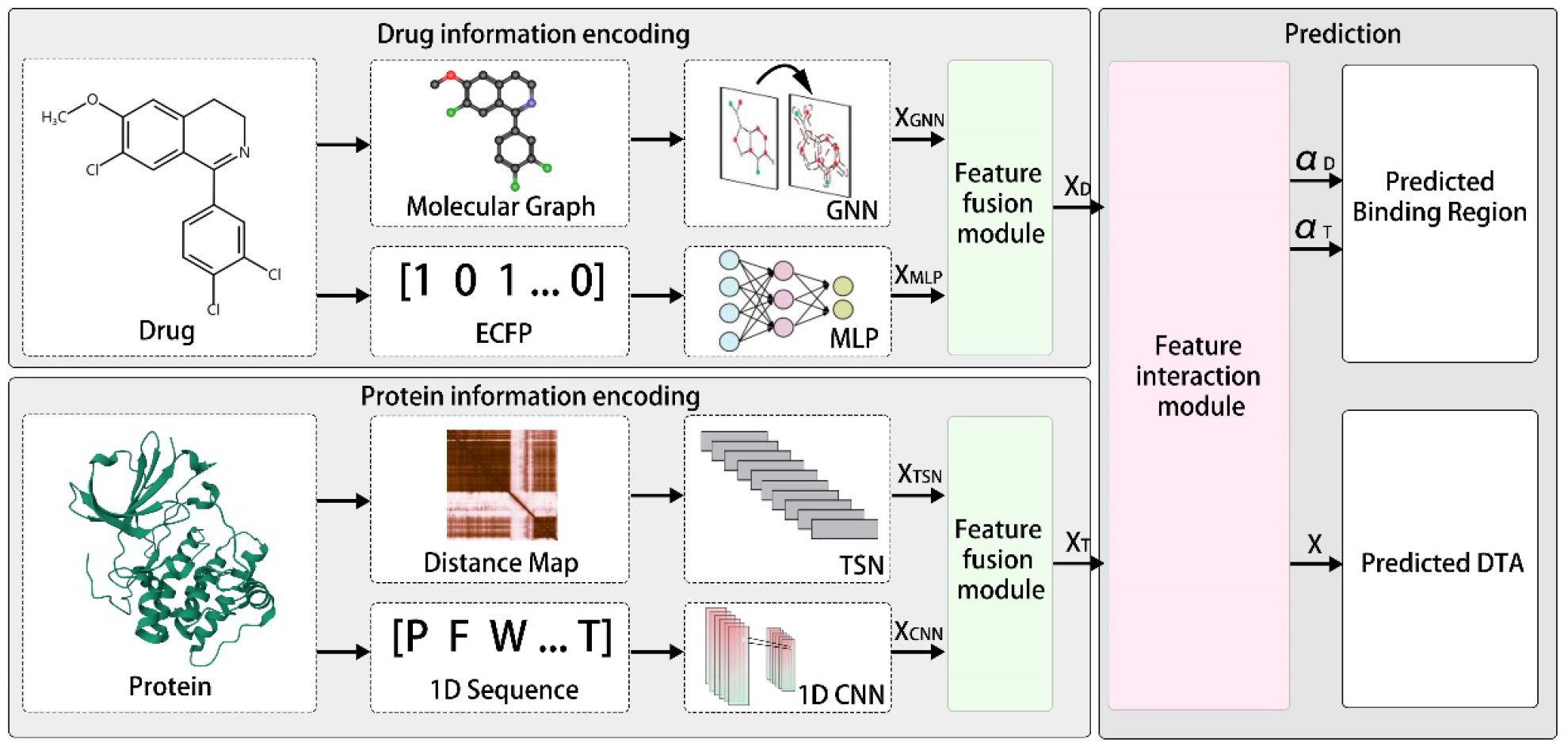
The workflow of MMD-DTA.

#### 2.2.1 Drug information encoding

Since molecular fingerprints contain a lot of useful physicochemical information, we choose to use extended connectivity fingerprints (ECFP) for drug molecular characterization [36]. Meanwhile, to obtain the structural information in the drug molecules, we construct molecular graph to further enrich the features extracted from ECFP. Each drug molecule is described as the SMILES sequence in the original data, and we use the open-source software RDkit to generate molecular fingerprint features and obtain information about adjacent inter-atomic bonds based on the SMILES sequence. Each atom is represented by a 78-dimensional embedding vector, encoded by the atomic elements (44 dimensions), the number of directly-bonded atoms (11 dimensions), the total number of H bound to the atom (11 dimensions), the total number of implicit H bound to the atom (11 dimensions), and whether the atom is aromatic (1 dimension). To ensure that the length of the features of different drugs is the same, we fixed the length of the drug sequence to *l=100*, and the shorter drugs are padded with 0. Finally, the fingerprint feature of the drug sequence is represented as *F*_*d*_ ∈ *R*^*l*^ ^×**E**^*_d_*, where *E*_*d*_ represents the dimension of the embedding vector of the atoms in the drug.

We use MLP model to process fingerprint feature *F*_*d*_. Due to the shorter length of drug sequences, MLP models with fewer linear layers can maintain their efficiency while preventing as much loss of individual information in drug atoms as possible. To avoid the negative impact of the shattered gradient problem [37] on model training, we use the skip connection structure [38] to connect the features between different layers. The process can be formulated as:

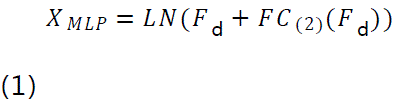

Where, *X*_*MLP*_ ∈ *R^*ld*×*Cd*^* is the output feature, *l*_*d*_ and *C*_*d*_ are the length and channel size of the drug fingerprint feature respectively, *LN*() represents a linear layer, *FC*_(2)_ is composed of two linear layers and a *ReLU* activation function.

GNN captures complex structural information by aggregating features between adjacent nodes and has been widely used in various fields of bioinformatics. Recently, GIN (Graph Isomorphism Network) proposed by xu *et al.* is known as the optimal graph representation learning method. We are inspired by GIN to process molecular graph *G*_*d*_(*V*,*E*) through an improved GNN network. The molecular graph *G*_*d*_(*V*,*E*) consists of node *V* (atom) and edge *E* (bond), and we aggregate the features between the relevant nodes as follows:

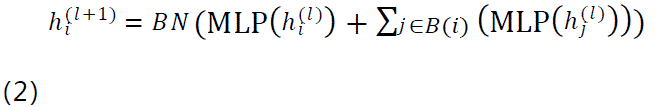

where, *BN* is the batch normalization layer, ℎ*_i_*^(*l*)^ and ℎ*^i^*^(*l*+1)^ are the feature representation of node *V*_*i*_ at layer *l* and layer *l+1* respectively, *MLP()* represents the multi-layer perceptron, and *B*(*i*) is the node set adjacent to node *V*_*i*_. Specifically, when *l=0*, we encode each node using the fingerprint feature as the original feature. After GNN processing, the structural features of the drug are expressed as *X*_*GNN*_.

#### 2.2.2 Protein information encoding

For protein sequences, we first represent each of its residues as specific initial features using one-hot encoding. Then by concatenating the embedding layers, each amino acid in the protein sequence is represented by a 128-dimensional vector. Finally, the original embedding of the protein sequence is represented as *F*_*t*_ ∈ *R*^*L*×*Et*^, where *E*_*t*_ represents the dimension of the embedding vector of the residues in the protein, and *L* is the length of the protein sequence, we fixed *L=1200*, and the shorter proteins are padded with 0. Next, we use 1D CNN to effectively learn abstract features from the original embeddings of protein sequences and retain individual information about each residue. Specifically, each convolutional layer consists of multiple convolution kernels, and different kernels act on different residue regions. For a given protein embedding feature *F*_t_, the calculation of the *n*th convolution kernel acting on the *i*th residue region is as follows:

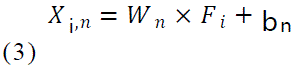

where, *X*_*i*,*n*_ ∈ *R*^*Lt*×*Ct*^ is the output feature; *L*_*t*_ and *C*_*t*_ are the length of the protein embedding feature and the channel size of the convolutional kernel respectively, *F*_*i*_ represents the embedding feature of the *i*th residue region of the protein, *W*_*n*_ and *b*_*n*_ represent the weight and bias of the *n*th convolutional kernel, respectively. After the convolution layer processing, the protein sequence feature is expressed as *X*_*CNN*_.

To obtain the structural information of proteins, we generate 2D pairwise distance maps by calculating the distances between each amino acid inside the target, and use the pairwise distance maps to reveal the true 3D structural information of proteins. For a protein sequence of length *L*, we use the simple and open-source PconsC4 model to predict the matrix *M*_t_ with *L* rows and *L* columns. The elements in row *i* and column *j* of the matrix correspond to the distance (Euclidean distance) between the *i*th amino acid and the *j*th amino acid in the protein.

After obtaining the 2D pairwise distance map of the target, we use the target structural feature extraction network (TSN) [39] composed of a large-kernel group convolutional layer and a self-attention module to effectively extract structural features and more global features in the protein. As shown in Figure 2, the network consists of a convolutional layer, a batch normalization layer, a linear layer, a *ReLU* activation function, and a *Softmax* activation function. The target structure feature extraction network can be formulated as:

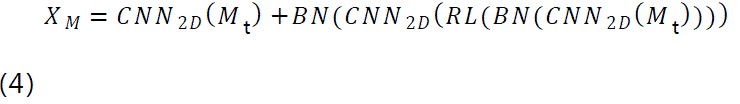

**Figure 2.**
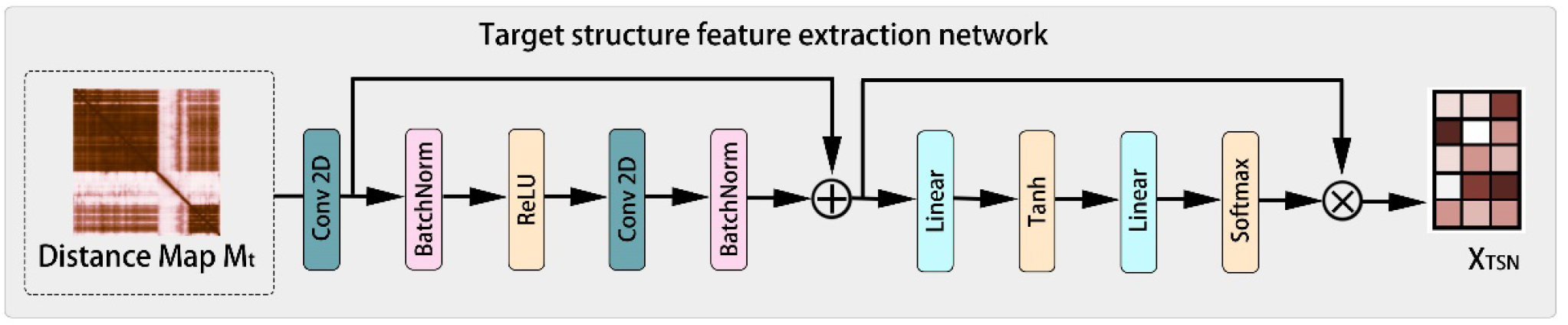
The structure of the target structure feature extraction network.

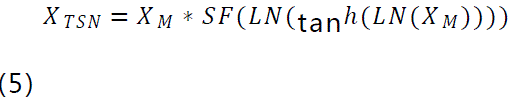

Where, *M*_*t*_ ∈ *R*^*L*×*L*^ and *X*_*TSN*_ ∈ *R*^*Lt*×*Ct*^ are the input and output of the target structure feature extraction network, *L*_*t*_ and *C*_*t*_ are the length of the protein embedding feature and the channel size of the convolution kernel respectively, *CNN*_2*D*_ represents 2D convolution block, *BN* is batch normalization layer, *LN* is linear layer, *RL* and *SF* represent *ReLU* activation function and *Softmax* activation function respectively. Equation (4) represents the process of the network capturing the spatial structure information between adjacent amino acids through the large-kernel group 2D CNN. Equation (5) is the self-attention module. Since the attention score can distinguish the importance of each feature channel, we give different weights to each amino acid in the protein to strengthen the important structural information and ignore the unimportant information. After TSN processing, protein structural features are represented as *X*_*TSN*_.

#### 2.2.3 Feature fusion module

As mentioned above, we obtain multiple features of drugs and targets through different feature extraction methods, which are drug fingerprint feature *X*_*MLP*_, drug structure feature *X*_*GNN*_, protein sequence feature *X*_*CNN*_, and protein structure feature *X*_*TSN*_. However, feature fusion is an important step in model construction and model performance improvement. To better learn the relationship between features and improve the expression ability of fusion features, we propose a new feature fusion module as shown in Figure 3. The calculation process of feature fusion is as follows:

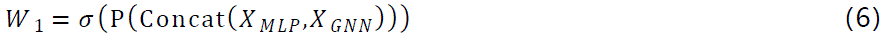

**Figure 3.**
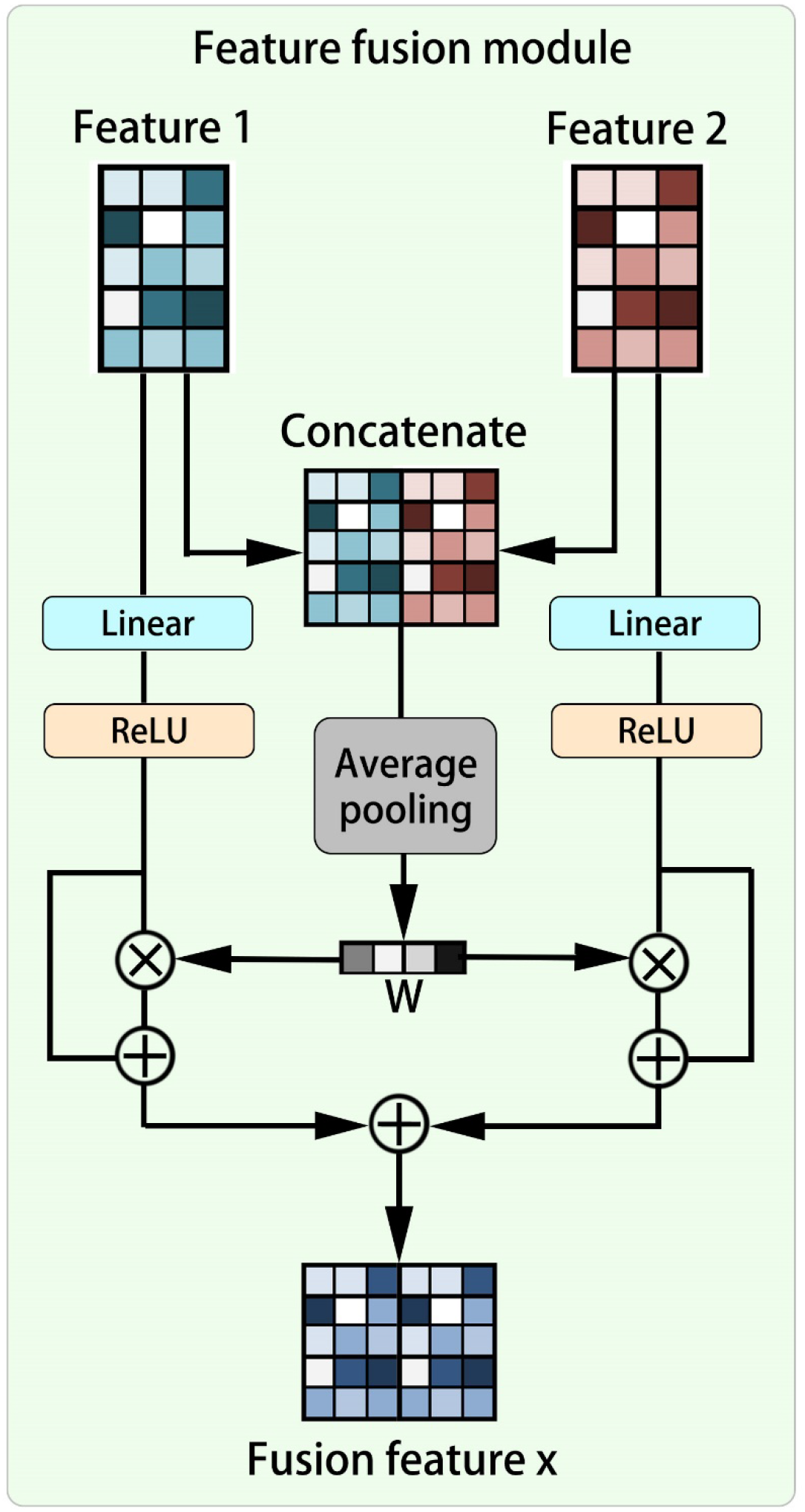
The structure of the feature fusion module.

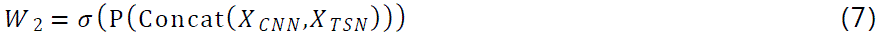

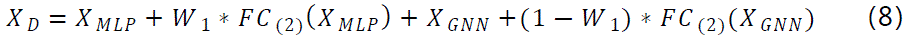

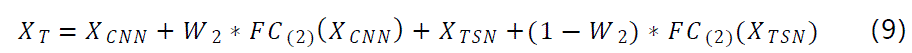

Where, *FC*_(2)_ is composed of two linear layers and a *ReLU* activation function, * is the element-wise multiplication, *σ* is the *Sigmoid* activation function, *P* is the global average-pooling function, *Concat* is the concatenation operation. Equations (6) and (7) represent the process of capturing the information of each site in the feature through the global average-pooling function and obtaining the feature attention weight *W* through the *Sigmoid* function. *(1-W)* represents the attention weight of the second feature to promote the balance of complementarity and contribution between different features. Finally, as shown in equations (8) and (9), we combine different features by residual connection to obtain the drug feature representation *X*_*D*_ and target feature representation *X*_*T*_.

#### 2.2.4 Feature interaction module

In this section, we design a feature interaction module as shown in Figure 4, which aims to obtain interaction features while performing drug-target BR prediction. We take the fusion feature *X*_*D*_ ∈ *R*^*ND*×*F*^ of the drug and the fusion feature *X*_*T*_ ∈ *R*^*NT*×*F*^ of the target as the input of the module, where *N*_*D*_ and *N*_*T*_ represent the sequence length of the drug and the target respectively, and *F* represents the depth. To predict the drug-target BR, we first obtain the drug feature kernel *D* and the protein feature kernel *T* through fully connected layer. Then, the attention score α ∈ *R*^*ND*×*NT*^ of each site is obtained through the tanh activation function. Finally, we obtain the interaction strength α_*D*_ ∈ *R*^*ND*^ of each site in the drug sequence to the target by applying row-wise sum operation to α, and the interaction strength α_*T*_ ∈ *R*^*NT*^ of each site in the target sequence to the drug by applying column-wise sum operation to α. The calculation process of the interaction strength of each site is as follows:

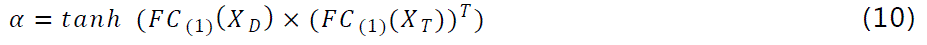

**Figure 4.**
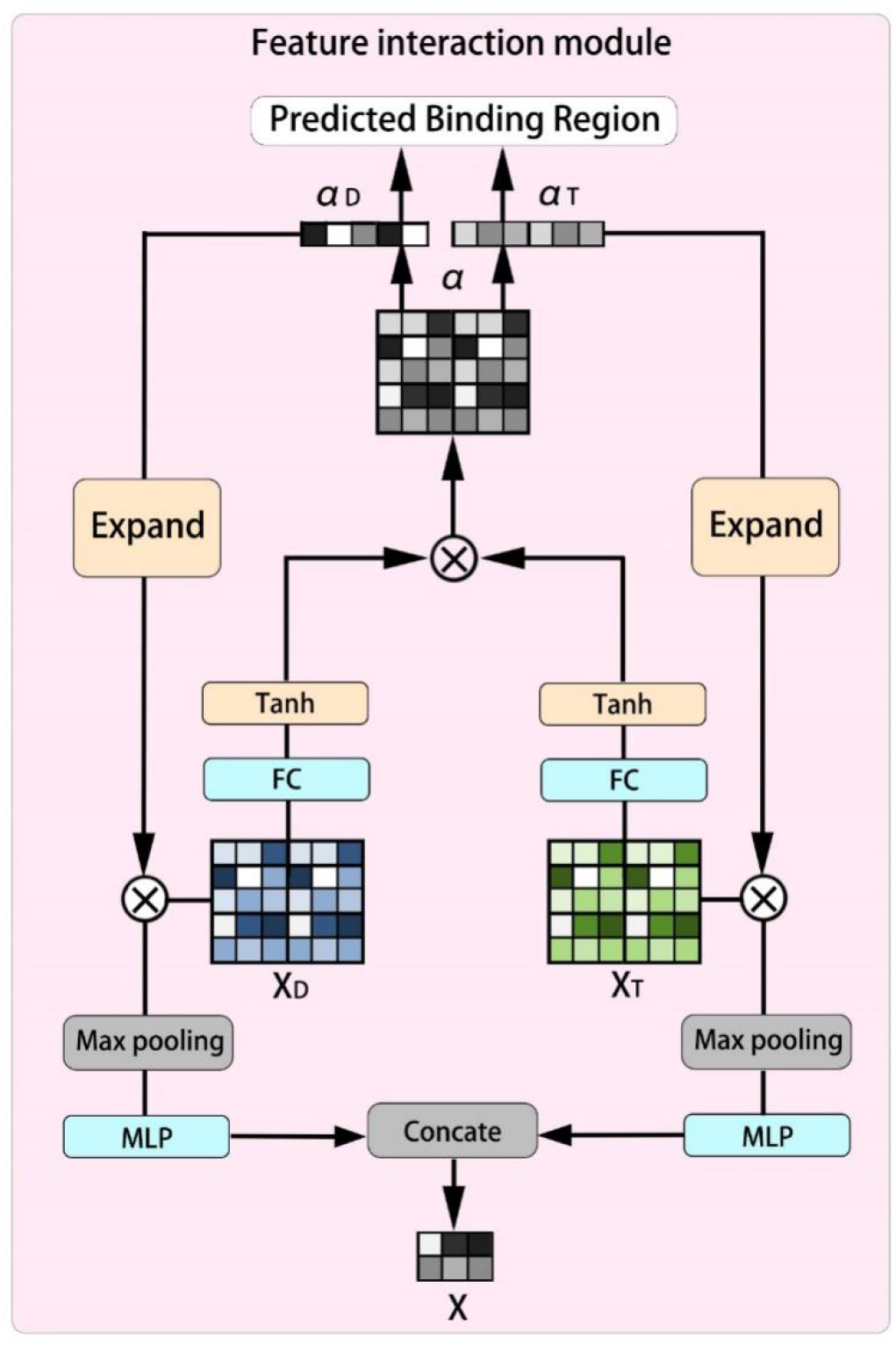
The structure of the feature interaction module.

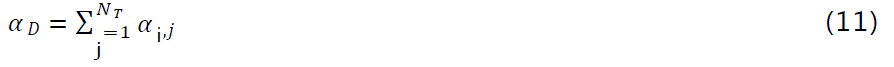

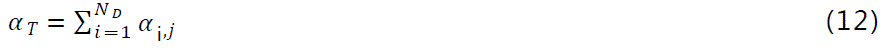

Where, *FC*_(1)_ is composed of the fully connected layer and *LeakyReLU* activation function, × is matrix multiplication and *T* is the matrix transpose. The binding affinity score α indicates the strength of the interaction between site in the drug sequence and site in the protein sequence.

In order to obtain the interaction drug features and interaction target features, we first expand α_*D*_ and α_*T*_, and obtain matrices *A*_*D*_ ∈ *R*^*ND*×*F*^ and *A*_*T*_ ∈ *R*^*NT*×*F*^ respectively by repeating the padding operation. Then, the feature representations of drugs and targets are enhanced by the *MLP* model and element-wise multiplication. Finally, we capture the most important feature information (the feature with the highest value) in the drug feature and target feature through max pooling function:

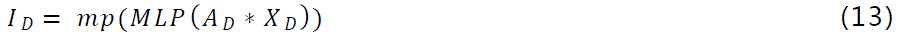

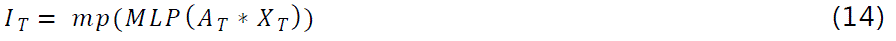

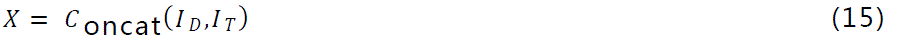

Where, *MLP()* represents multi-layer perceptron, *mp* is the max pooling function, *\** is the element-wise multiplication, *I*_*D*_ ∈ *R*^*F*^ and *I*_*T*_ ∈ *R*^*F*^ are the final feature representations of drugs and targets respectively. After the feature interaction processing, we connect *I*_*D*_ and *I*_*T*_ to obtain the drug-target interaction descriptor *X* ∈ *R*^2*F*^.

### 2.3 Prediction and training

We input the drug-target descriptors into an FCNN consisting of 3-layer neural networks. A dropout layer is attached behind both the first and second layers to prevent the model from overfitting. Finally, we get the predicted value of DTA from the output of the third layer and the prediction of the BR from the interaction strength of each site in the target sequence.

We use MSE to measure the gap between the DTA predicted value *P* and the real value *Y* and use it as the loss function of the model. The equation is as follows:

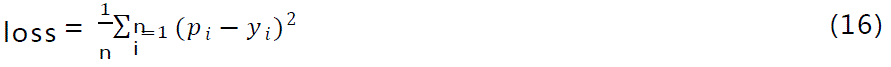

Where *p*_*i*_ and *y*_*i*_ represent the predicted and real values of the affinity of the *i*th drug-target pair, respectively.

The model in this paper is implemented in Python3.7.16 with Torch1.4.0. We further divided the dataset into training dataset and test dataset by 5-fold CV, using Adam optimizer for parameter training. Table 2 shows the details of hyper-parameter Settings.

**Table 1.**
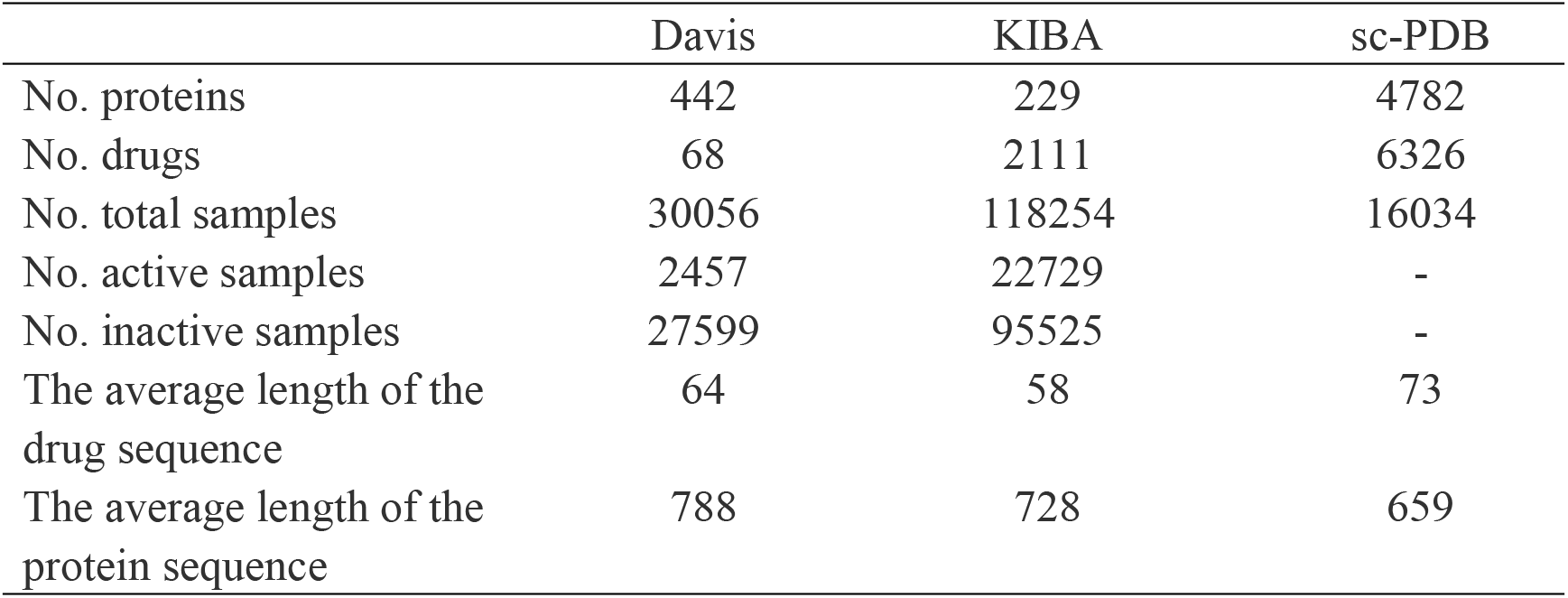
Summary of the benchmarking datasets.

|  | Davis | KIBA | sc-PDB |
| --- | --- | --- | --- |
| No. proteins | 442 | 229 | 4782 |
| No. drugs | 68 | 2111 | 6326 |
| No. total samples | 30056 | 118254 | 16034 |
| No. active samples | 2457 | 22729 | - |
| No. inactive samples | 27599 | 95525 | - |
| The average length of the drug sequence | 64 | 58 | 73 |
| The average length of the protein sequence | 788 | 728 | 659 |

**Table 2.** Hyper-parameter settings of our model.

| Hyper-parameters | Setting |
| --- | --- |
| Learning rate | 0.0005 |
| Batch size | 512 |
| GNN layers | 3 |
| Kernel size of $CNN_{1D}$ | 8 |
| Kernel size of $CNN_{2D}$ | (1200, 1) |
| $E_d, E_t$ | 78, 128 |
| $N_D, N_T, F$ | 128, 128, 128 |
| $l_d, C_d, L_t, C_t$ | 128, 32, 128, 32 |

## 3. Results

### 3.1 Evaluation metrics

Consistent with previous evaluation metrics for DTA prediction problems, we use MSE and Concordance Index (CI) [40] as evaluation metrics to evaluate the performance of the proposed model. CI mainly reflects whether the predicted value and the real value of the binding affinity of the drug-target pair are in the same order (the larger the value, the more accurate the model prediction), which is calculated by equation (8):

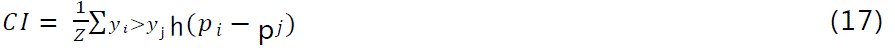

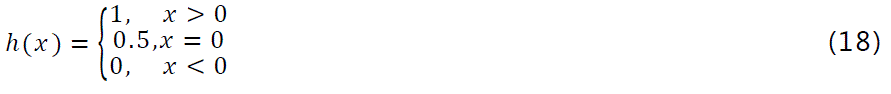

where *Z* is a normalization constant and *h (x)* is a step function.

MSE is the mean of the sum of squared differences between the predicted value *P*_i_(*i* = 1,2,⋯,*n*) and the real value *Y*_i_(*i* = 1,2,⋯,*n*) (the smaller the value, the more accurate the model prediction), calculated by equation (19):

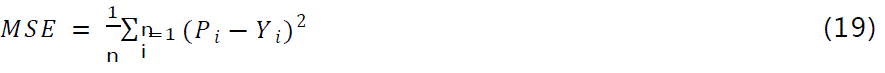

### 3.2 Comparison with the state-of-the-art methods

In this section, we use 5-fold CV to evaluate the predictive performance of MMD-DTA on the Davis and KIBA datasets. At the same time, we compare MMD-DTA with the six current state-of-the-art DTA prediction models to demonstrate the superiority of our model. The baseline model is described as follows:

- DeepDTA is a DTA prediction algorithm based on the convolutional layer and the fully connected layer [9].
- CPInformer performs CPI prediction by extracting compound molecular graph features, fingerprint information, and multi-scale protein features [15].
- DeepCDA uses the LSTM algorithm and CNN to predict the compound protein affinity [21].
- MATT-DTI is a prediction method for DTI with a multi-head attention module [41].
- GraphDTA extracts the structure information of drug molecules through GNN and combines protein features to predict DTA [42].
- DeepGLSTM uses the Bi-LSTM algorithm and GCN to extract protein sequence features and drug features, respectively, to predict the binding affinity of drug targets [43].

Table 3 shows the performance of MMD-DTA and the baseline model, respectively. On the Davis dataset, MMD-DTA achieves the highest CI value (0.905 on average) and the lowest MSE value (0.220 on average). Compared with the best baseline approach (DeepGLSTM), MMD-DTA achieves a performance gain of 1.3% and 6.7% in terms of CI and MSE, respectively. Compared to GraphDTA, MMD-DTA achieves a performance gain of 1.7% and 5.6% in terms of CI and MSE, respectively, mainly because our proposed protein coding module provides more extensive and fine protein features for the model, thus improving the performance of the model.

**Table 3.**
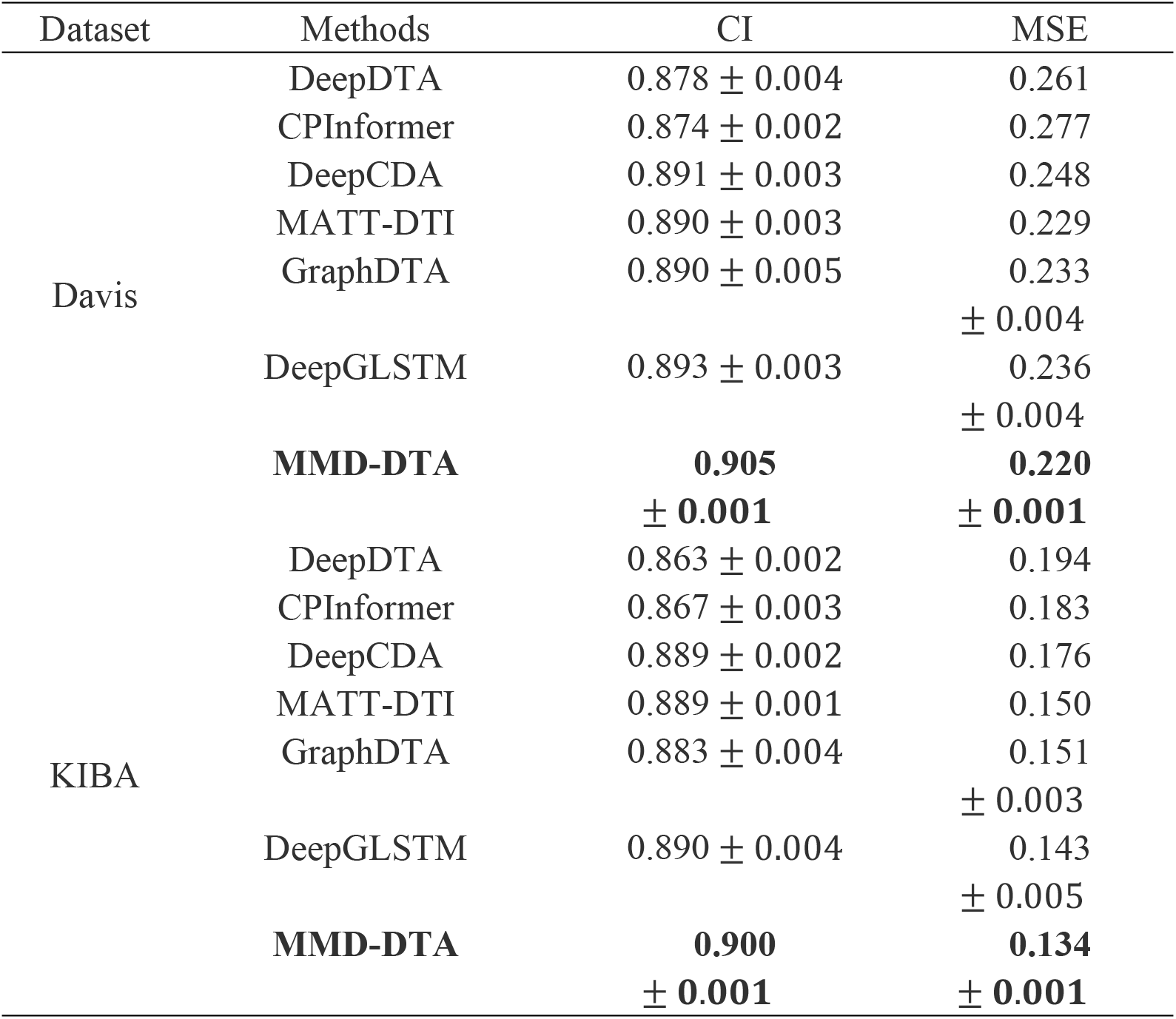
Performance comparison of MMD-DTA and state-of-the-art methods on the Davis and KIBA datasets, bold: best results.

On the KIBA dataset, the optimal CI value obtained by MMD-DTA is 0.900. It is 4.3%, 3.8%, 1.2%, 1.2%, 1.9%, and 1.1% higher than DeepDTA, CPInformer, DeepCDA, MATT-DTI, GraphDTA, and DeepGLSTM, respectively. The average MSE value of MMD-DTA is 0.134, which is 30.9%, 26.8%, 23.9%, 10.7%, 11.2%, and 6.3% lower than DeepDTA, CPInformer, DeepCDA, MATT-DTI, GraphDTA, and DeepGLSTM, respectively. However, the CI values of almost all models on the KIBA dataset are lower than those on the Davis dataset. This may be due to the concentration of sample labels in the KIBA dataset, which makes it difficult for most models to predict trends in binding affinity. In general, MMD-DTA achieves more accurate prediction results than the baseline model on both Davis and KIBA datasets.

### 3.3 External validation

To evaluate the generalizability of the model, we reset the training and test datasets on the Davis dataset so that there is no overlap between the training and test datasets and neither the training drugs nor training targets appear in the test datasets. The performance comparison between MMD-DTA and the baseline model under 5-fold CV on the newly set Davis dataset is shown in Figure 5. The results show that MMD-DTA performs well on the newly set Davis dataset. For unseen drugs and targets, MMD-DTA achieves the highest CI value of 0.733, which is 37.5%, 34.3%, 49.6%, 59.7%, 41.8%, and 14.9% higher than DeepDTA, CPInformer, DeepCDA, MATT-DTI, GraphDTA, and DeepGLSTM, respectively. The MSE value of MMD-DTA is 0.413, which is 34.5%, 31.0%, 24.8%, 51.2%, 38.5%, and 5.3% lower than DeepDTA, CPInformer, DeepCDA, MATT-DTI, GraphDTA, and DeepGLSTM, respectively.

**Figure 5.**
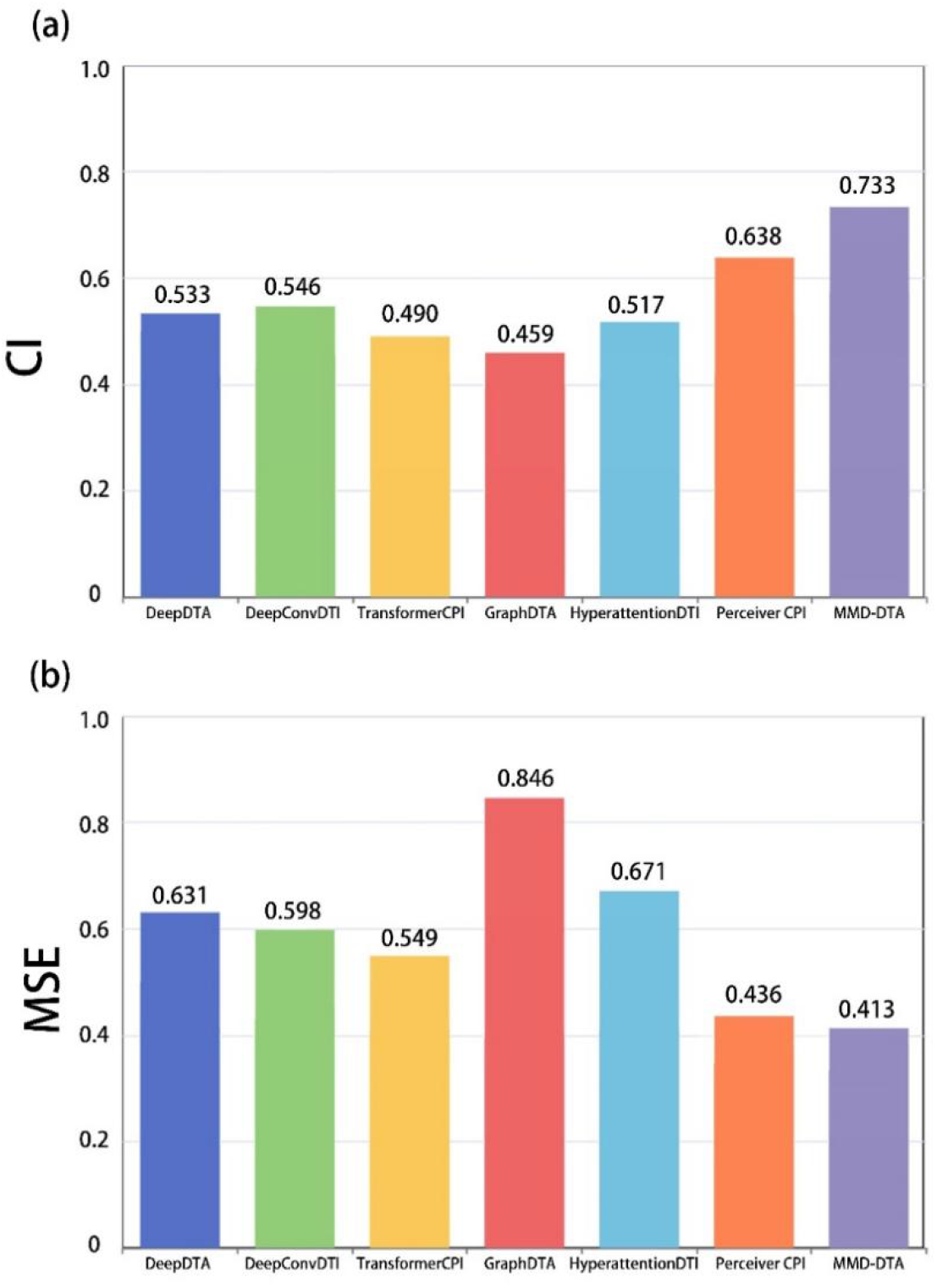
(a) CI results of MMD-DTA and baseline models on the newly set Davis dataset. (b) MSE results of MMD-DTA and baseline models on the newly set Davis dataset.

In addition, to evaluate the adaptability of MMD-DTA to new domain datasets, we conduct another cross-domain experiment. Specifically, we first eliminate all parts of the independent test dataset (PDBbind) that overlap with the benchmark dataset (Davis). Then, we train the model on the Davis dataset and test it on the PDBbind dataset. The cross-domain experimental results of MMD-DTA and other models are shown in Figure 6. The results show that MMD-DTA is significantly better than the baseline model. Compared with other models, MMD-DTA obtains the best CI value of 0.573, which is 14.6%, 20.0%, 15.3%, 11.0%, 39.8%, and 7.8% higher than DeepDTA, CPInformer, DeepCDA, MATT-DTI, GraphDTA, and DeepGLSTM, respectively. The MSE value of MMD-DTA is 4.271, which is 9.4%, 20.9%, 13.9%, 32.5%, 28.2%, and 7.4% lower than DeepDTA, CPInformer, DeepCDA, MATT-DTI, GraphDTA, and DeepGLSTM, respectively.

**Figure 6.**
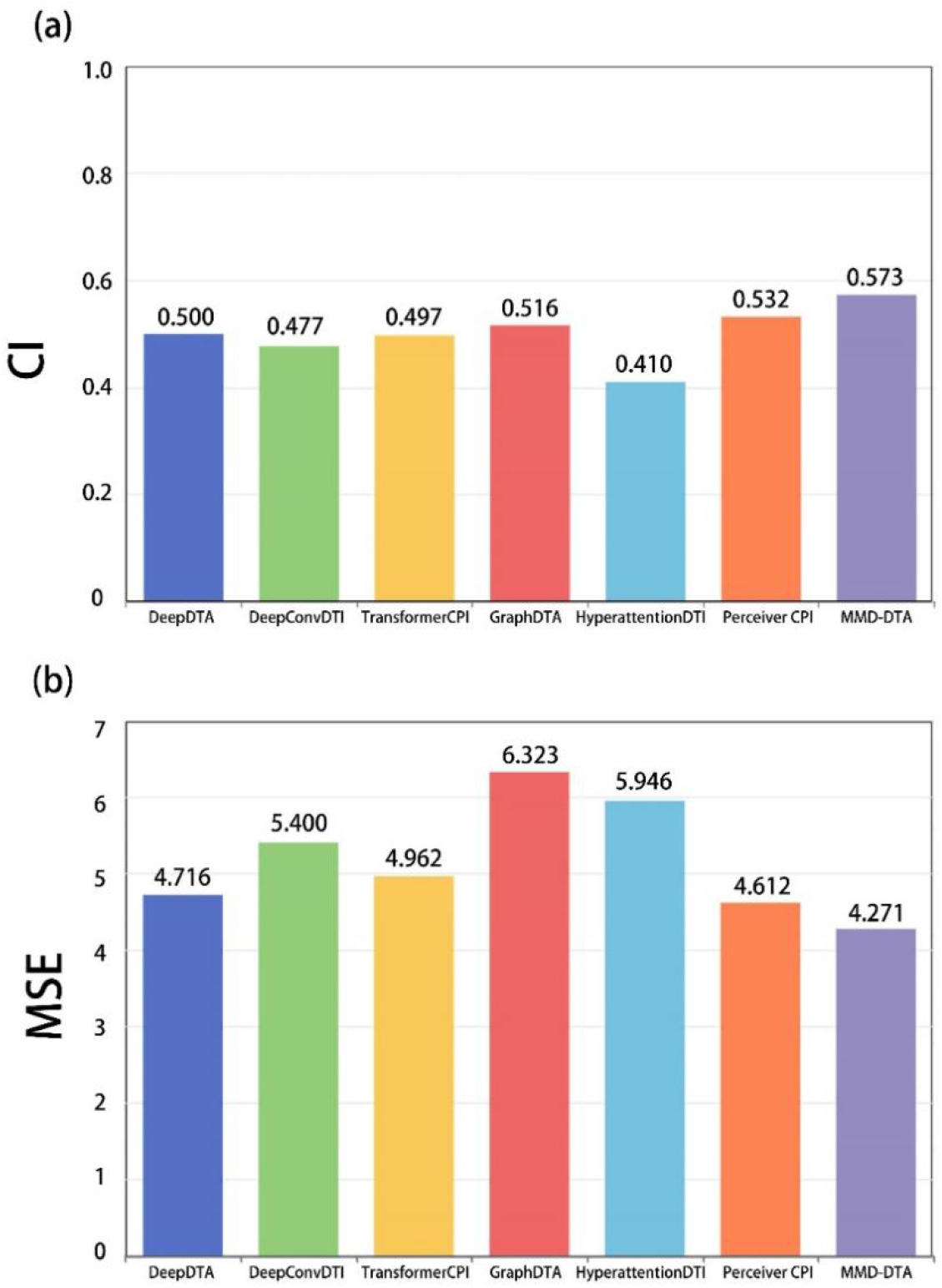
(a) CI results of MMD-DTA and baseline models on cross-domain datasets. (b) MSE results of MMD-DTA and baseline models on cross-domain dataset.

In general, MMD-DTA achieves more accurate prediction than the baseline model on the newly set Davis dataset and on the cross-domain dataset. This shows that the model trained on the benchmark dataset can generalize well to practical applications. However, based on the results of external validation, we observed a performance gap between drugs and targets seen and unseen. MMD-DTA extracts comprehensive features including drug fingerprint features, drug structural features, protein sequence features and structural features. More comprehensive features help us improve the accuracy of the unseen DTA prediction, but there is still a lot of room for improvement.

### 3.4 Ablation study

The innovative elements of MMD-DTA include sequence information encoding, structural information encoding, feature fusion, feature interaction, and other key modules. In this study, we evaluate the effectiveness of each innovative element of the model and the importance of different drug and protein extraction methods. To evaluate the effect of each component on the model performance, we conduct ablation studies on Davis and KIBA datasets using 5-fold CV.

- Model-1: We remove the fingerprint features in the drug information encoding and the sequence features in the protein information encoding, and we directly input the remaining features into the feature interaction module.
- Model-2: We remove the structural features in the drug information encoding and protein information encoding and directly input the remaining features into the feature interaction module.
- Model-3: We remove the feature attention weights *W*_1_ and *W*_2_ in the feature fusion module, and we remove the residual connection and directly connect the features.
- Model-4: We directly connect the output *X*_D_ and *X*_T_ of the feature fusion module to remove all operations of the feature interaction module.

Table 4 shows a comparison of the CI and MSE performance between the variants and MMD-DTA. The table shows that the optimal MSE value obtained by MMD-DTA on the Davis dataset is 0.220, which is 25.7% and 27.4% lower than Model-1 and Model-2, respectively. MMD-DTA achieves the highest CI value of 0.905, which is 5.6% and 4.9% higher than the above two models, respectively. These results show that drug fingerprint features, structural features, protein sequence features, and structural features all contain useful information, and MMD-DTA extracts more comprehensive features (sequence features and structural features), thus improving the prediction performance of the model. Compared to Model-3, MMD-DTA has a 1.7% improvement in CI and a 6.4% reduction in MSE on the Davis dataset. This verifies the superiority of the proposed feature fusion module. Compared to Model-4 on the Davis dataset, the CI of MMD-DTA increases by 0.9% and the MSE decreases by 5.2%. The results show that the feature interaction module can not only predict the drug-target BR, but also plays a crucial role in the prediction of DTA. At the same time, MMD-DTA also has different levels of performance improvement compared with various variants on the KIBA dataset. In summary, our innovative elements help the model to achieve the best results on both datasets.

**Table 4.**
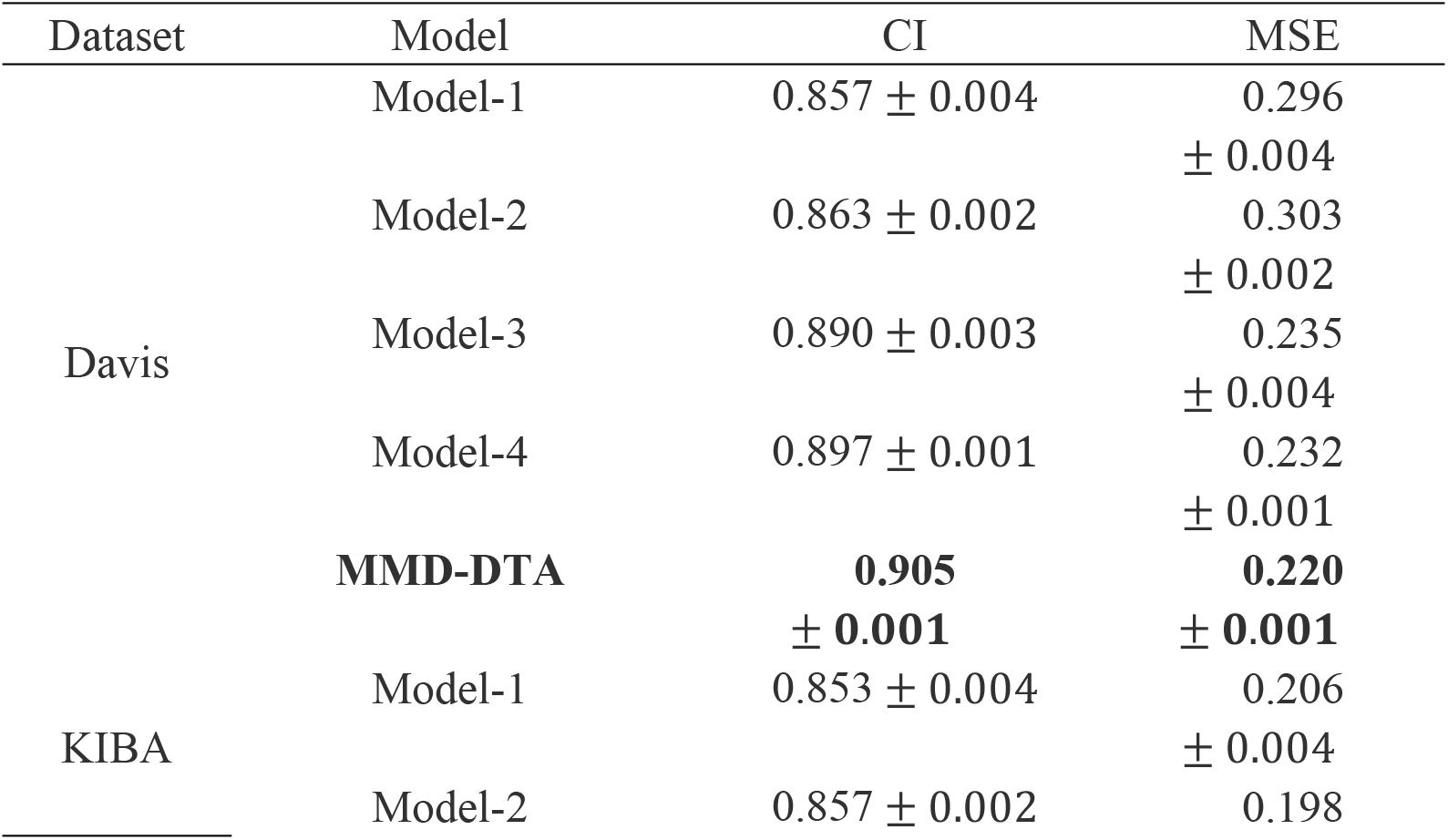
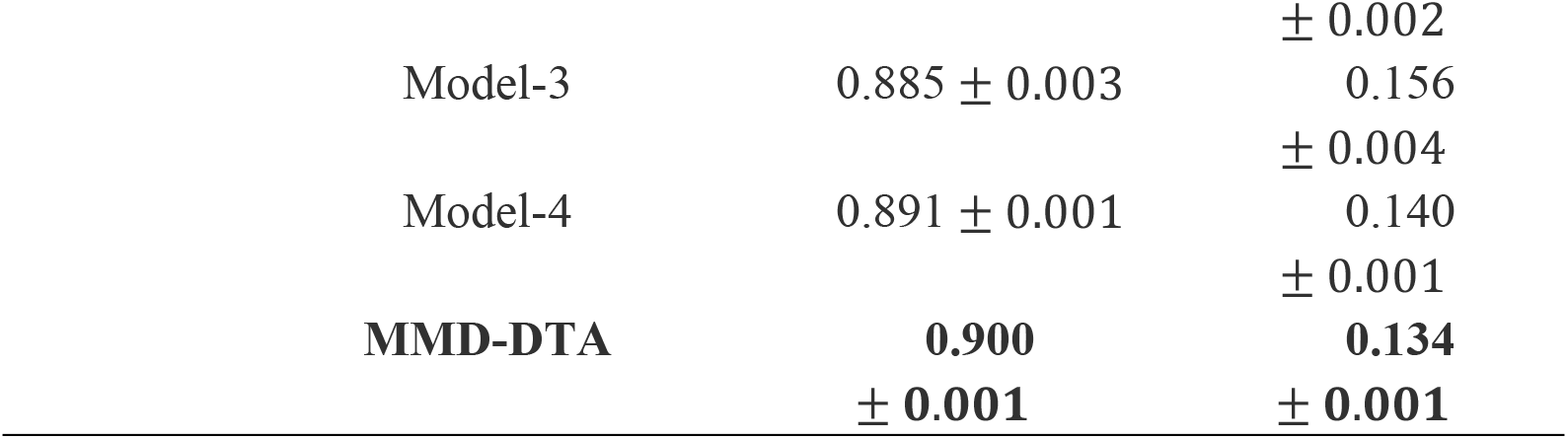
Results of Ablation Experiments, bold: best results.

| Dataset | Model | CI | MSE |
| --- | --- | --- | --- |
| Davis | Model-1 | $0.857 \pm 0.004$ | $0.296 \pm 0.004$ |
| | Model-2 | $0.863 \pm 0.002$ | $0.303 \pm 0.002$ |
| | Model-3 | $0.890 \pm 0.003$ | $0.235 \pm 0.004$ |
| | Model-4 | $0.897 \pm 0.001$ | $0.232 \pm 0.001$ |
|  | **MMD-DTA** | **0.905** | **0.220** |
|  |  | **$\pm 0.001$** | **$\pm 0.001$** |
| KIBA | Model-1 | $0.853 \pm 0.004$ | $0.206 \pm 0.004$ |
| | Model-2 | $0.857 \pm 0.002$ | $0.198$ |

|  |  |  |
| --- | --- | --- |
| | | $\pm 0.002$ |
| Model-3 | $0.885 \pm 0.003$ | 0.156 |
| | | $\pm 0.004$ |
| Model-4 | $0.891 \pm 0.001$ | 0.140 |
| | | $\pm 0.001$ |
| **MMD-DTA** | **0.900** | **0.134** |
|  | **$\pm 0.001$** | **$\pm 0.001$** |

### 3.5 Drug–target BR prediction

To show the predictive power of MMD-DTA for drug-target BR, we compare MMD-DTA with existing unsupervised approaches to predict drug-target BR. Specifically, we first select the site with the highest value from the interaction strength of each site of the target sequence obtained in Section 2.5. Then, the protein sequence fragment with the length of 10 amino acids is cut forward and backward with the highest value site as the center, and this sequence fragment is used as the prediction region of drug-target binding. Finally, we take the probability of the actual binding site falling into the predicted region as a metric to measure the accuracy of the model.

Table 5 shows the prediction effects of different models on the BR on Davies, KIBA, and sc-PDB datasets. On the Davis dataset, when the scale of the prediction region is 10, the prediction accuracy of MMD-DTA is 0.322, which is better than other methods. When we apply the model trained on the Davis dataset to the sc-PDB dataset, MMD-DTA achieves the best prediction accuracy (0.307), which is 22.6%, 11.6%, 7.0%, and 9.4% higher than CPI-GNN, DeepCDA, TransformerCPI, and CPInformer, respectively. On the KIBA dataset, the prediction accuracy of MMD-DTA (0.505) is also higher than that of other methods, and when we apply the trained model to the sc-PDB dataset, its accuracy reaches 0.430, which is 33.7%, 27.9%, 23.4%, and 21.2% higher than CPI-GNN, DeepCDA, TransformerCPI, and CPInformer, respectively.

**Table 5.** Comparison of the performance of MMD-DTA and different methods in predicting the drug-target BR, bold: best results.

| Dataset | Methods | S=10 | sc-PDB |
| --- | --- | --- | --- |
| Davis | CPI-GNN | 0.126 | 0.081 |
|  | DeepCDA | 0.257 | 0.191 |
|  | TransformerCPI | 0.213 | 0.237 |
|  | CPInformer | 0.158 | 0.213 |
|  | **MMD-DTA** | **0.322** | **0.307** |
| KIBA | CPI-GNN | 0.313 | 0.093 |
|  | DeepCDA | 0.472 | 0.151 |
|  | TransformerCPI | 0.374 | 0.196 |
|  | CPInformer | 0.488 | 0.218 |
|  | **MMD-DTA** | **0.505** | **0.430** |

We demonstrate the interpretability of the model by visualizing the prediction of the BR of the test sample. Here, Figure 7 (a) shows the predicted BR of drug “CHEMBL1087421” and protein “O00141”, and Figure 7 (b) shows the predicted BR of drug “CHEMBL208637” and protein “O14757”. Where the regions marked in red are true binding sites but predicted by the model as not binding sites, the regions marked in black are binding sites accurately predicted by the model, and the regions marked in blue are not binding sites but predicted by the model as binding sites. Obviously, the binding site of protein “O00141” completely falls into the predicted region of the model, which intuitively shows the excellent performance of MMD-DTA in predicting the drug-target BR. Although the prediction of the binding site of the protein “O14757” is somewhat biased, a large part of the predicted region overlaps with the binding site. At the same time, we also visualize the prediction region of the binding site of the drug “CHEMBL1087421” and the prediction region of the binding site of the drug “CHEMBL208637”. This visualization has important biomedical significance, and exploring the functional regions of drug molecules acting on proteins is an important idea to enhance the biomedical interpretability of the model.

**Figure 7.**
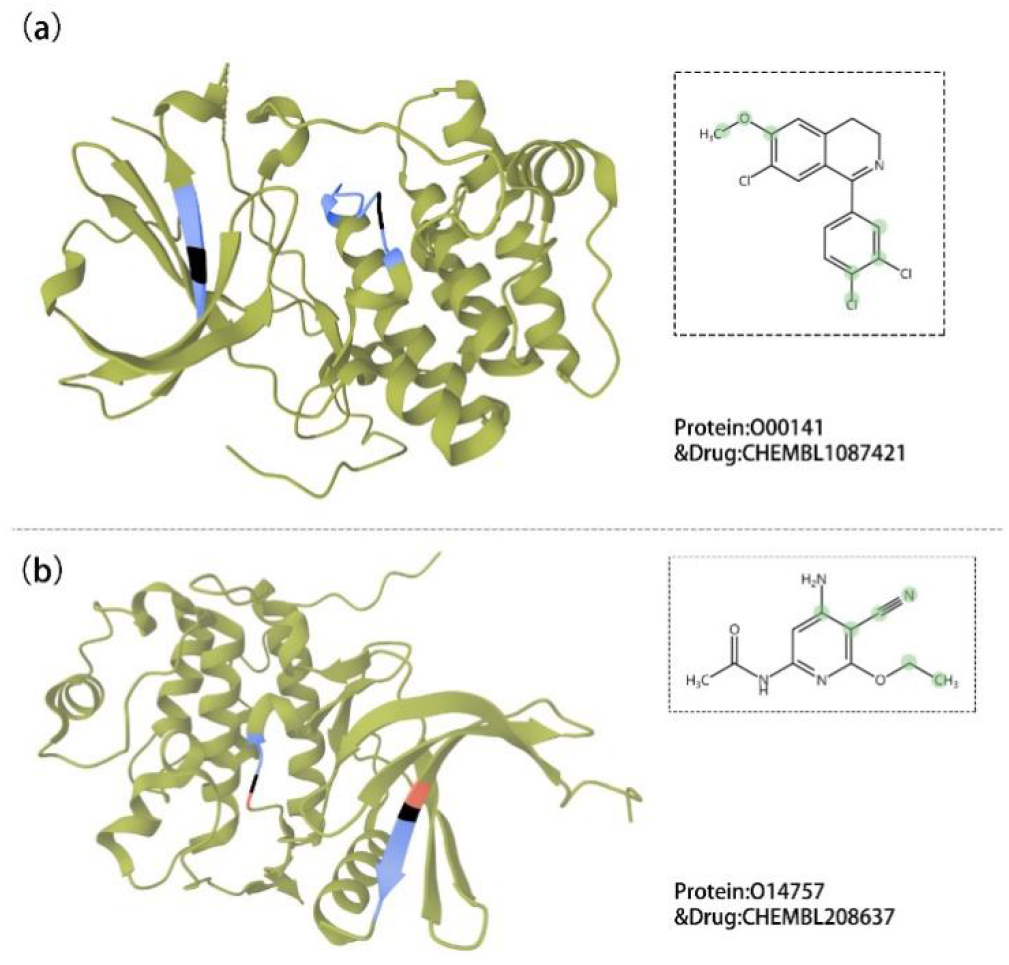
(a) Visualization of BRs for the drug “CHEMBL1087421” and protein “O00141”. (b) Visualization of BRs for the drug “ CHEMBL208637” and protein “O14757”.

## 4. Discussion and conclusion

In recent years, deep learning has developed rapidly, and more and more deep learning models have been applied to DTA prediction. Although previous DTA prediction methods have achieved good results, how to simply and effectively obtain more information about DTA and predict drug-target BR is still a difficult problem to be solved. In this work, we construct a multi-modal deep learning framework (MMD-DTA) for drug-target binding affinity and BR prediction. We demonstrate the superiority of our model by comparing MMD-DTA with six current state-of-the-art DTA prediction models. In order to demonstrate the interpretability of MMD-DTA and its ability to predict drug-target BR, we visually analyze the prediction results of BR. In general, MMD-DTA is an effective model for predicting drug-target binding affinity and BR.

There are four main innovative modules that improve the predictive performance of MMD-DTA. First, MMD-DTA improves GIN and MLP and further utilizes the structural information of drug molecules while obtaining the physicochemical information of drug molecules. Second, we extract the sequence information of the protein from the amino acid sequence using 1D CNN, and extract the structural information of the protein from the protein distance map constructed by the PconsC4 model by using the target structural feature extraction network. Third, the feature fusion module helps the model to better learn the relationship between features, improve the expression ability of features, and achieve the balance of complementarity and contribution between different features. Fourth, the feature interaction module can simultaneously output the drug-target descriptor for DTA prediction and the interaction strength of drug sequence and target sequence at each site for drug-target BR prediction, which enhances the interpretability of the model and provides important information for further drug structure optimization.

However, MMD-DTA still has some room for improvement. First, the protein distance map is generated by the PconsC4 model. In recent years, the methods to represent the 3D structure information of proteins have been improved, and there may be more suitable methods to represent the structural characteristics of proteins. Second, the hyperparameters of MMD-DTA on the Davis and KIBA datasets are the best hyperparameters selected by us after referring to the hyperparameter Settings of the previous methods [46], and there may be better hyperparameters to improve the prediction performance of MMD-DTA. In the future, we will explore more efficient and rapid protein structure characterization methods and try to find better hyperparameters to further improve the prediction accuracy of MMD-DTA.

## Data availability

The datasets used in this study and the source code for MMD-DTA are publicly available at https://github.com/wyx2012/MMD-DTA.

## Acknowledgments

This work was supported by the open research fund of Key Laboratory of Computational Science and Application of Hainan Province (No. JSKX202102,JSKX202301).

## Appendix A

CI and MSE results of MMD-DTA and baseline models on the newly set Davis dataset. (Table A1); and CI and MSE results of MMD-DTA and baseline models on cross-domain datasets. (Table A2).

**Table A1.**
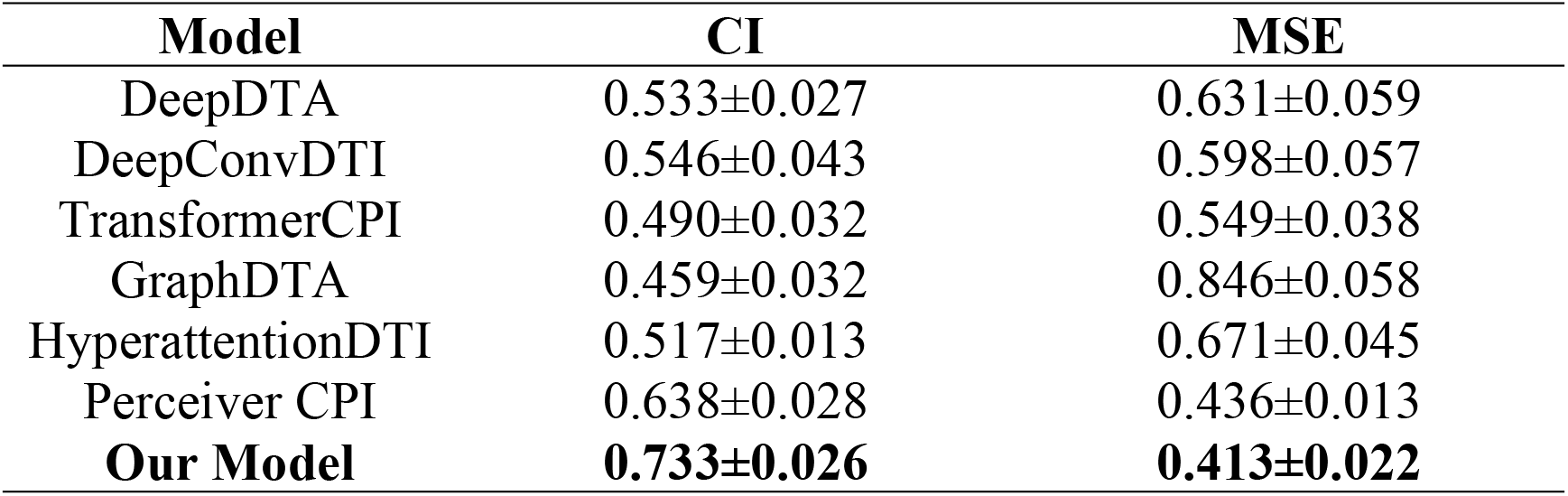
CI and MSE results of MMD-DTA and baseline models on the newly set Davis dataset.

| Model | CI | MSE |
| --- | --- | --- |
| DeepDTA | 0.533±0.027 | 0.631±0.059 |
| DeepConvDTI | 0.546±0.043 | 0.598±0.057 |
| TransformerCPI | 0.490±0.032 | 0.549±0.038 |
| GraphDTA | 0.459±0.032 | 0.846±0.058 |
| HyperattentionDTI | 0.517±0.013 | 0.671±0.045 |
| Perceiver CPI | 0.638±0.028 | 0.436±0.013 |
| **Our Model** | **0.733±0.026** | **0.413±0.022** |

**Table A2.** CI and MSE results of MMD-DTA and baseline models on cross-domain datasets.

| Model | CI | MSE |
| --- | --- | --- |
| DeepDTA | 0.500±0.027 | 4.716±0.596 |
| DeepConvDTI | 0.477±0.043 | 5.400±0.572 |
| TransformerCPI | 0.497±0.032 | 4.962±0.387 |
| GraphDTA | 0.516±0.032 | 6.323±0.582 |
| HyperattentionDTI | 0.410±0.013 | 5.946±0.450 |
| Perceiver CPI | 0.532±0.028 | 4.612±0.131 |
| **Our Model** | **0.573±0.026** | **4.271±0.22** |

## List of abbreviations

DTI: drug-target interaction
DTA: drug–target affinity
BR: binding region
FCNN: fully connected neural network
GCN: graph convolutional network
CNN: convolutional neural network
5-fold CV: 5-fold cross validation
CI: concordance index
MSE: mean squared error

